# Beaver Impacts Tool: A Graphical User Interface for Analyzing Ecosystem Impacts of Beaver Dams Using Remote Sensing Data

**DOI:** 10.64898/2026.11.12.766214

**Authors:** Quintin Tyree, Jack Etheredge, Ziyu Kong, Cary Murray, Chris Norcross, Wendy A Owens-Rios, Grace W Lindsay

## Abstract

Beavers are being increasingly recognized as driving factors in diverse and resilient riparian ecosystems. Many of their impacts, including changes to water levels and vegetation, can be studied through remote sensing. However, ecologists and conservationists may lack the tools to easily analyze remote sensing data for these purposes. We offer here a Google Earth Engine-backed Streamlit app that allows users to easily upload locations of observed dams or installed beaver dam analogs for analysis. Via a graphical user interface, users then generate a set of dam-absent locations for comparison and receive plots and numerical data indicating how indices associated with vegetation, water, temperature, and evapotranspiration change over time for dam-present and dam-absent locations. Based on the availability of open datasets, our tool provides coverage from 2017 onward. The cloud version of the tool can be accessed via a static URL, or users can run the app locally using code and documentation provided on github. We believe this tool will allow a wider range of ecologists and conservationists to complement their work with the use of remote sensing data, without the need to gain expertise in specific software.

## 1 INTRODUCTION

The North American beaver (*Castor canadensis*) is a keystone species and ecosystem engineer known for its ability to transform landscapes by altering hydrology, biogeochemistry, and plant communities in the habitats it occupies (Naiman, Johnston, & Kelley, 1988; Johnston & Naiman, 1990; Naiman, Melillo, & Hobbie, 1986; Naiman, Pinay, Johnston, & Pastor, 1994; Fairfax & Westbrook, 2024). By constructing dams and creating surface-water impoundments, beavers expand wetlands, increase habitat heterogeneity, and create a mosaic of aquatic, emergent, and riparian habitats (Naiman et al., 1994; Brazier et al., 2021; Grudzinski et al., 2022). Historically widespread across North America, beavers were nearly extirpated by the early 20th century primarily due to over-trapping and extensive habitat loss, including the draining of wetlands and development along riparian corridors (Zedler, 2004; Mitsch, Gosselink, Zhang, & Anderson, 2023).

The use of beavers as a tool for ecosystem restoration has gained increasing attention because of their ability to instigate ecological processes through their natural engineering activities. Beaver reintroduction and translocation can provide a form of process-based restoration, allowing beavers to modify hydrology, increase habitat complexity, and promote the development of wetland and riparian ecosystems (Law, Gaywood, Jones, Ramsay, & Willby, 2017; Doden, Budy, Conner, & Young, 2023). In addition, beaver dam analogs (BDAs) are increasingly used to mimic the effects of natural beaver dams in degraded stream and riparian systems where beavers are absent or unable to establish (Wohl, 2021). Despite the growing recognition of beavers as a nature-based restoration tool, beaver-related restoration remains relatively underrepresented in the scientific literature, accounting for only 4.3% of the beaver-management literature reviewed by Treves and Comino (Treves & Comino, 2024). This relatively limited representation highlights an opportunity to better understand both the ecological benefits of beaver-driven restoration and the challenges associated with incorporating beavers into restoration and management strategies.

In addition to the scientific study of beaver impacts, momentum is building to better integrate beavers into restoration and climate resilience strategies. But this requires inclusive, data-informed frameworks (Nash et al., 2021). Recent work emphasizes the importance of combining ecological theory with practical tools to improve restoration outcomes and ensure ecological knowledge translates into action (Silliman et al., 2024). Spatial decision-support tools can further assist restoration practitioners and land managers by, for example, identifying where beaver activity may provide restoration benefits and where potential conflicts may occur (Macfarlane et al., 2017). However, access to specialized tools, data, and technical expertise can limit the broader application of beaver-based approaches and research. Developing accessible tools that translate ecological information into usable knowledge may therefore expand practitioners’ ability to incorporate beavers into restoration planning and provide evidence of their impacts.

Publicly available satellite imagery provides an additional opportunity to support ecological monitoring by allowing changes in ecosystems to be evaluated consistently across broad spatial and temporal scales (Pettorelli et al., 2014, 2018). Tools that integrate these data can streamline assessments and generate standardized, spatially explicit information across sites. Such information can assist researchers and land managers in tracking ecological change and evaluating restoration outcomes. This provides evidence that can inform management decisions, guide policy, and enhance ecological understanding (Sutherland, Pullin, Dolman, & Knight, 2004; Pullin & Knight, 2009)

Here, we introduce the Beaver Impacts Tool, a Streamlit app with a Google Earth Engine (GEE) backend, that assesses impacts of beaver dams using satellite imagery. The app provides a user-friendly graphical interface with step-by-step instructions, resulting in informative plots and downloadable data. This tool can be accessed by anyone online, or run locally using a user’s GEE API key.

## 2 MATERIALS AND METHODS

With this tool, we aim to make analysis of remote sensing data for beaver impacts as simple as possible. With our graphical user interface (GUI), users start by providing coordinates of dam locations, e.g. like those identified in a dam census. From there, they are presented with options on how to proceed based on their data needs or restrictions. An overview and example of the process is shown in Fig. 1. The output of the tool is a visualization and optional download (in CSV format) of monthly values capturing vegetation, water, temperature, and evapotranspiration metrics at dam-present and to dam-negative locations, for user-specified year(s). This allows users to observe any impacts of dams on these metrics. Below we explain the technical details of the tool implementation.

**FIGURE 1.**
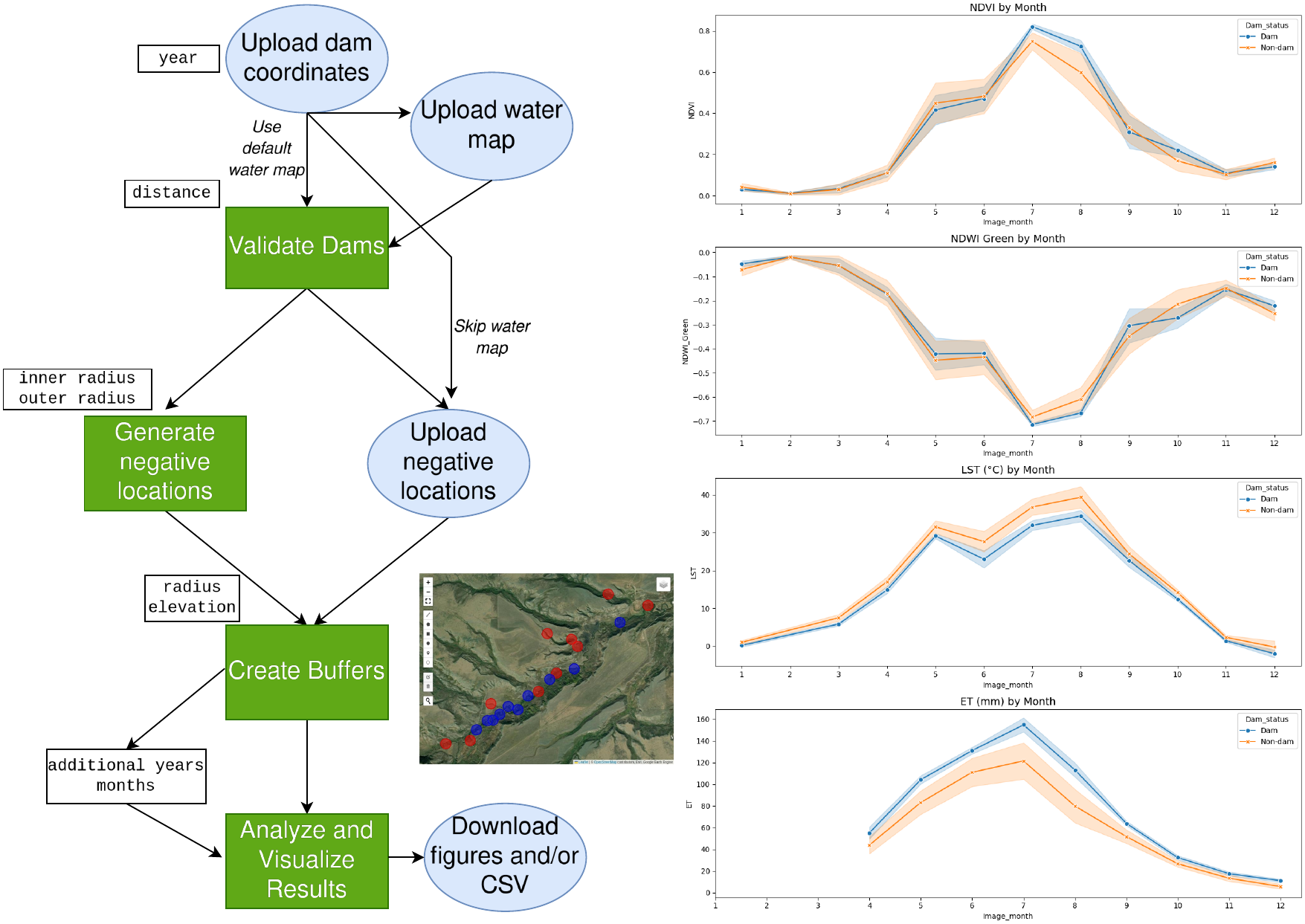
Flow diagram of steps in the tool. Blue circles indicate user uploads, green squares are performed by the tool, white squares indicate user-defined variables. Inset image shows what the tool-generated map looks like at the buffer stage. On the right are the results plotted by the tool (OpenET data was missing for some months, which the tool indicates to the user before plotting).

**FIGURE 2.**
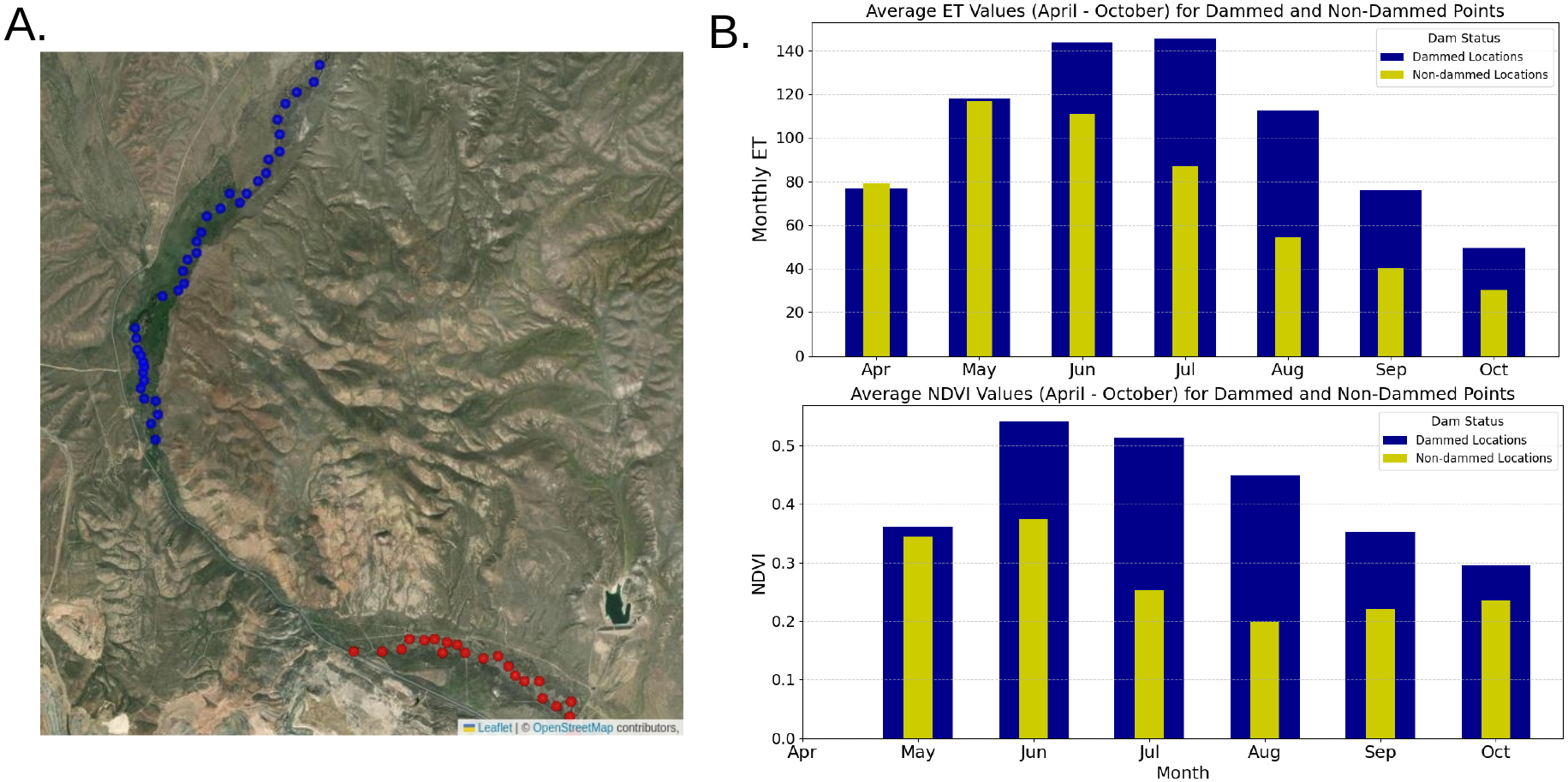
Replication of Maggie Creek results from (Fairfax & Small, 2018). A.) Points generated to represent the identified dammed (blue) and non-dammed (red) regions. B.) Monthly ET (top) and NDVI (bottom) values for dammed and non-dammed regions in 2017 (comparable to Fig. 10 in (Fairfax & Small, 2018)). Note: acceptable Sentinel-2 imagery was not available for April in this location.

### 2.1 Implementation

The tool is implemented as a Streamlit web application in Python using the geemap library to leverage GEE’s geospatial processing capabilities. A flow diagram of tool functions and user inputs can be seen in Fig. 1.

#### 2.1.1 Dam Location Upload and Validation

The analysis begins with the upload of a dataset containing known beaver dam locations (referred to as ‘dam-positive’ sites) in either CSV or geoJSON format. CSV files should contain a latitude and a longitude column. Users also select a year for the analysis (following imagery data availability, years can be from 2017 onward). These sites are then stored as a GEE FeatureCollection (e.g., “projects/lab/assets/DamData”).

The dataset is then validated to ensure that dam points are located on or near waterways. By default, waterway data are sourced from the National Hydrography Dataset (NHD) available in GEE (Moore et al., 2019). The study area is defined by the bounding box of the positive dam locations and U.S. state boundaries (TIGER/2018/States) are filtered to identify states intersecting this area. For each state, the corresponding NHD dataset is retrieved and merged into a single FeatureCollection, creating a comprehensive waterway dataset for analysis. In addition to NHD, users also have the option of using an alternative waterway map developed by the World Wildlife Foundation, called “WWF HydroSHEDS Free Flowing Rivers Network v1” (Lehner, Verdin, & Jarvis, 2008). Users may also upload their own waterway map directly in geoJSON format or through GEE. For each dam feature, a buffer is created around its geometry, and nearby waterways are identified. The distance method computes the shortest distance to any waterway feature. Dams within the maximum allowed distance (default 50m) are classified as valid, while those exceeding it are flagged as invalid. Following user agreement, the validated dam locations are used as the positive dam dataset for subsequent steps, ensuring that only geographically accurate points are analyzed.

If waterway maps are not available (e.g. if the coordinates are outside the coverage area of NHD and/or WWF), users can choose to skip dam validation and automatic dam-negative site generation (which rely on waterway data) and instead move directly to uploading their own dam-negative sites.

#### 2.1.2 Negative Site Generation

To form a comparison group, locations without dams (‘dam-negative sites’) can be uploaded by the user (CSV or geoJSON), or be automatically-generated. For the latter option, negative points are generated to be along waterways but away from known dams. Note that this process requires that all dams in the analyzed region have been identified and uploaded as positive sites; otherwise, negative sites may be unknowingly chosen near unlabeled dams.

Negative points are sampled within a ring-shaped region created by buffering positive locations by an inner radius (default 300 meters) and an outer radius (default 500 meters), then taking the difference. Using the default values, this results in dam-negative selection regions that are greater than 300m but less than 500m away from any known dam. The stratified Sample tool samples points within this ring that are near waterways. Where possible, the number of negative sites chosen will be equal to the number of dam-positive sites, ensuring a balanced dataset for comparative analysis.

#### 2.1.3 Buffering and Sample Region Definition

Sample regions are created around both dam-positive and dam-negative locations to define areas for data extraction. To do so, each dam point (positive or negative) is buffered by a (default) 150-meter radius to create circular sample regions. Elevation data from the USGS 3DEP 10m dataset is used to refine sample regions (similar to a procedure used in (Fairfax & Small, 2018)). Specifically, the elevation at each dam point was sampled, and a mask was created to include only points in the buffer region that were within -5 to +3 (default) meters of the dam’s elevation. This ensured the sample regions were focused on ecologically relevant areas near the waterway. In the case that elevation data is not available (e.g. outside the US), imagery data will be collected and used from the entire circular sample region.

#### 2.1.4 Remote Sensing Data Products

Sentinel-2 Surface Reflectance data (Drusch et al., 2012) were used to derive Normalized Difference Vegetation Index (NDVI) and Normalized Difference Water Index (NDWI). Images were filtered by date (±6 months from July 1 of the user-defined year(s)) and classified into calendar months. If multiple images were available within a single month, the median image was taken. Binary cloud masks were derived from the Sentinel-2 QA60 band when available to exclude cloudy and cirrus-affected pixels from the monthly median composites. NDVI was calculated as (NIR - Red) / (NIR + Red) and NDWI as (Green - NIR) / (Green + NIR). Mean values were computed over the valid pixels within each sample region to create a single value per month per location (positive or negative).

Land Surface Temperature (LST) was derived from Landsat 8 Collection 2 Level 2 data using established methodology (Malakar et al., 2018). If no images were available within a calender month, NaN values were assigned; if there were multiple images the median value was used. For each sample region, the LST band was clipped to the sample region and the mean was computed.

Evapotranspiration (ET) data were sourced from the OpenET Ensemble dataset (Melton et al., 2022). OpenET provides monthly ET values. These values are available for a subset of the locations and years that Sentinel-2 data is available, therefore not all queries will be able to return ET data. The exact availability of ET data varies as OpenET expands its coverage. If no data were available for a given month, NaN was assigned. Where available, the mean ET was computed for each sample region.

Data processing is divided into batches to manage computational load. Processing time is a function of the number of dams, combined with variability from backend resource fluctuations in Streamlit and GEE.

#### 2.1.5 Visualization and Download

Data processing progress is displayed to the user via a widget. Once completed, the extracted metrics (NDVI, NDWI, LST, ET) for dam-positive and dam-negative sites were converted to a Pandas DataFrame that is exportable as a CSV. The DataFrame was sorted by month and year, and invalid values were filtered out. Users have the option to measure impacts within a single year or across years. If plotting data for a single year, Seaborn line plots are generated to visualize values per month (shaded regions represent 95% confidence intervals), with separate subplots for each metric, comparing positive vs. negative sites (see Fig. 1 for an example).

If plotting trends across years, users indicate which months they want included in the annual average. Bar plots show mean values over selected months with 95% confidence error bars. See Fig. 3B for an example NDVI plot across years.

**FIGURE 3.**
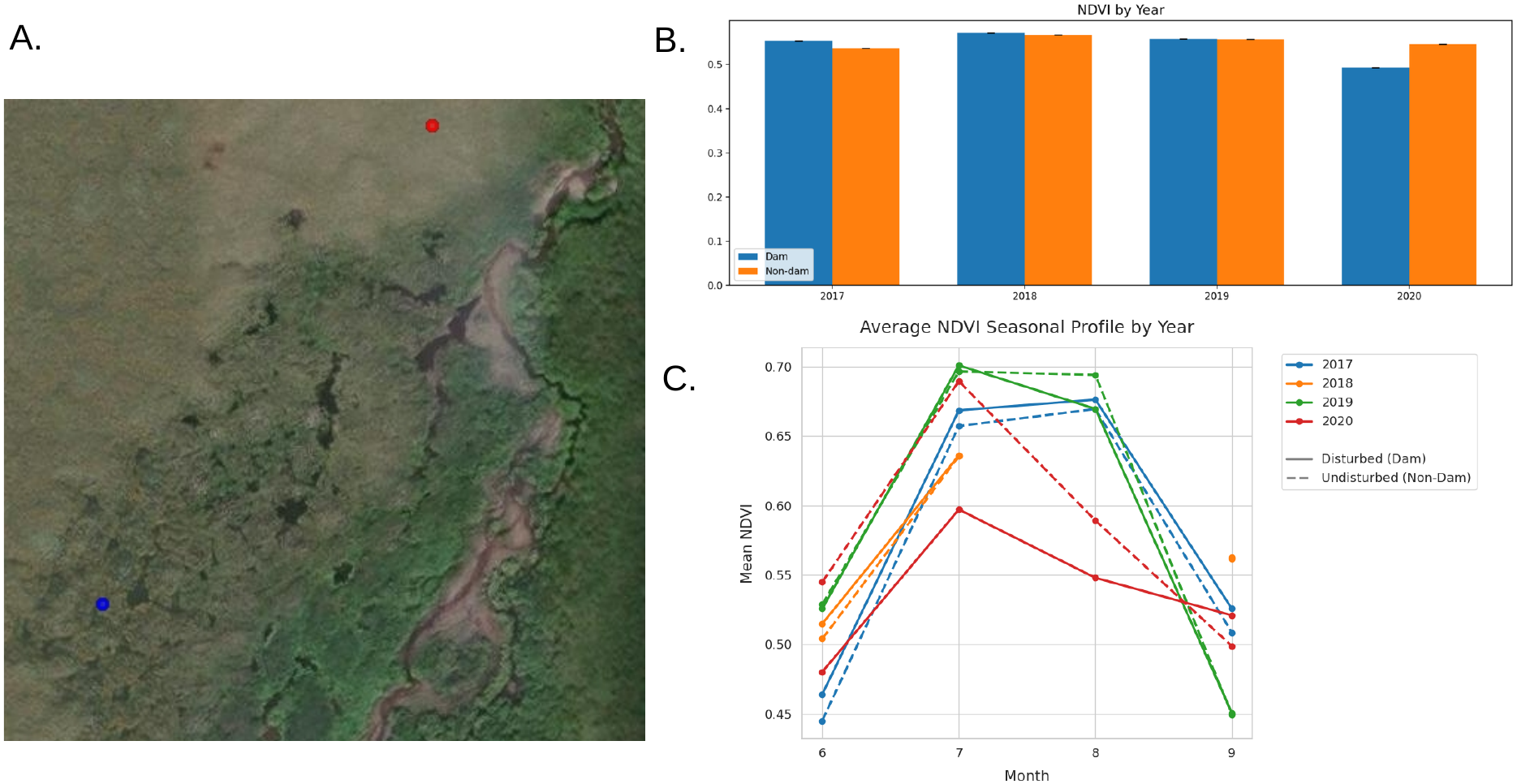
Replication of results from (Jones et al., 2021) using multi-year analysis. A.) Points analyzed to mimic the dam (blue) and control (red) locations from the study. B.) Output figure generated using the tool’s multi-year analysis. C.) Data plotted similarly to (Jones et al., 2021)’s Fig. 8 showing a change in the NDVI at the disturbed dam location in 2020. Note: insufficient imagery was available for August of 2018.

## 3 RESULTS

To demonstrate a typical experience with the tool, we apply it to a small set of dam locations in Fig. 1. The blue circles in the inset image indicate the circular buffer regions around the dam-positive locations. The red regions are the automatically generated dam-negative locations. The output of the analysis, shown on the right, indicates a moderate effect of dams on ET and temperature in the summer months, with a less robust impact on NDVI or NDWI.

### 3.1 Benchmarking and Validation

To demonstrate the validity of our data pipeline, we replicate previous results analyzing the impacts of dams in Maggie Creek, NV ((Fairfax & Small, 2018)). In this study, precise dam locations were not given, but dammed and undammed regions were compared. To replicate this, we generated a series of points in each of the previously analyzed regions (see Fig. 2A). We upload the points from the dammed regions as positive locations and the points from the undammed region as negative locations (using 3m elevation masks and 100m buffers). This previous study uses Landsat 8 images from 2013-2016, however, the Sentinel-2 data used in our tool is only available from 2017 onward so that is the year we use (note that due to a lack of suitable Sentinel imagery we do not have NDVI for April). Similar to Fairfax and Small, we observe consistently higher NDVI and ET in the dammed versus non-dammed regions for the months June through October (Fig. 2B). Fairfax and Small also sees these differences in May and, for ET, in April. The difference in our results is likely due to the difference in year, as 2013-2016 included drought conditions in NV, which likely amplified the impact of dams, whereas 2017 did not.

We also validate the tool’s ability to compare data across years by replicating an analysis from Jones et al. (Fig. 3). That study documented a period of heavy rainfall in August 2019 that caused an established dam system in Swan Lake Creek, AK to fail. The authors observed a decrease in NDVI at the disturbed location in 2020 compared to its previous values and compared to a control location. Using the same dammed and control regions as that paper (Fig. 3A, 50m buffers), we replicate the 2020 effect (Fig. 3B and C).

## 4 DISCUSSION

We created the Beaver Impacts Tool to allow researchers and practitioners from a wide range of backgrounds to easily access and benefit from freely-available remote sensing data. The tool lets users compare measures of vegetation, water, and temperature at dammed locations to those of nearby un-dammed locations in order to document the ecological impacts of beaver dams. We expect this tool to be useful to ecology researchers working in academia, in addition to land owners, conservationists, and policy makers outside academia. For example, researchers can use the tool on large-scale beaver dam census data to document beaver impacts across an entire state. Alternatively, a land manager who installed beaver dam analogs can use the tool to measure the impact on their property.

The tool currently has limitations, and we hope to offer more features in the future. For example, the algorithm for automatic dam-negative site selection chooses sites primarily based on their distance from known dams. It does not take into account other features of the waterway that may contribute to ecological differences (such as stream width). For a more tightly-controlled comparison, users should hand-select and upload negative locations. In the future, we would also like to be able to provide more impact metrics such as plant diversity or soil carbon content, however these depend on the state of technologies capable of deriving these measures from publicly-available satellite imagery ((Wood, Pidgeon, Radeloff, & Keuler, 2012; Gould, 2000; Reddy, 2021; Rasel, Groen, Hussin, & Diti, 2017; Nur et al., 2025)).

## DATA AVAILABILITY STATEMENT

We did not collect data for this study. Coordinates used to replicate past work are included in code files.

## AUTHORS’ CONTRIBUTIONS

HT, JE, ZK, and CM contributed to the software design and coding of the tool. CN and WAO contributed to the scientific design. GWL contributed to project supervision, conception, and design.

## ACKNOWLEDGEMENTS

ZK was supported by an NYU Research Catalyst Grant.

## ETHICS STATEMENT

No approvals required.

